# Metabolome genome-wide association study reveals hierarchical and epistatic genetic control of flavonoid metabolism in soybean

**DOI:** 10.64898/2026.07.23.739694

**Authors:** Taisei Hatta, Kosuke Hamazaki, Yushiro Fuji, Yusuke Toda, Yasunori Ichihashi, Yoshihiro Ohmori, Yuji Yamasaki, Hirokazu Takahashi, Hideki Takanashi, Mai Tsuda, Hisashi Tsujimoto, Akito Kaga, Mikio Nakazono, Toru Fujiwara, Masami Yokota Hirai, Hiroyoshi Iwata

## Abstract

Metabolic phenotypes are often governed by complex genetic architectures involving both additive and non-additive effects. However, the extent to which epistatic interactions contribute to the pathway-level regulation of plant metabolism remains unclear. In this study, we investigated the genetic architecture of flavonoid-related metabolites using metabolomic and genomic data from 200 soybean accessions cultivated under multiple environmental conditions. Broad-sense heritability estimates revealed that many metabolites were under strong genetic control, particularly flavonoid-related metabolites. Principal component analysis-based metabolome-wide genome-wide association studies identified four major loci associated with flavonoid metabolic variation, including a locus corresponding to flavonoid 3′-hydroxylase. Conditional analyses based on multilocus genetic backgrounds demonstrated that the effects of downstream loci were highly dependent on upstream genotypes. In particular, single-nucleotide polymorphism effects were frequently detectable only in specific allelic backgrounds defined by the major flavonoid 3′-hydroxylase locus, consistent with strong epistatic interactions among loci. Bayesian network analyses further supported a hierarchical genetic structure consistent with upstream regulation of downstream loci across the flavonoid biosynthetic pathway. These results demonstrate that highly heritable metabolic phenotypes can be controlled by a few loci exhibiting both additive and context-dependent non-additive effects. Our findings provide evidence that pathway-level metabolic diversity in soybean is generated through hierarchical and epistatic genetic control involving a limited set of key loci.

## Introduction

Plants produce a wide range of metabolites essential for growth, development, and environmental responses (Smith and Stitt, 2007; Saito and Matsuda, 2010). Among them, secondary metabolites such as flavonoids play key roles in plant defense and adaptation (Wink, 2003; Hartmann, 2007). Considering that the accumulation of these metabolites is a quantitative trait influenced by both genetic and environmental factors, elucidating the genetic basis of metabolic variation is important for improving crop functionality and metabolite production (The Arabidopsis Genome Initiative, 2000; Zabala and Vodkin, 2003; Butelli et al., 2008; Fernie and Schauer, 2009; Naqvi et al., 2009; Stitt et al., 2010; Litvinov et al., 2021).

Genome-wide association studies (GWAS) have been widely used to investigate the genetic architecture of such traits (Yu et al., 2006). Specifically, metabolome-wide GWAS enable the simultaneous analysis of multiple metabolites, facilitating the identification of genomic regions with pleiotropic effects and providing insights into the genetic control within complex metabolic networks (Riedelsheimer et al., 2012; Matsuda et al., 2015).

However, metabolic pathways are organized as coordinated and hierarchical systems, in which downstream reactions depend on upstream processes. Thus, genetic effects at downstream loci may only be expressed under specific genetic backgrounds, particularly when upstream reactions are active. This leads to context-dependent genetic effects, commonly referred to as epistasis (Lehner, 2011; Soltis and Kliebenstein, 2015). In metabolic pathways, such context-dependent effects may arise because downstream genetic effects often depend on the activity of upstream reactions. As a result, observed interactions may reflect biochemical dependencies rather than purely statistical associations (Rowe et al., 2008; Soltis and Kliebenstein, 2015). Although epistasis is recognized as an important component of complex trait variation (Carlborg and Haley, 2004; Phillips, 2008; Moore and Williams, 2009; Saito and Matsuda, 2010; Mackay, 2014), it is not adequately captured by standard GWAS models that primarily estimate additive effects at individual loci.

In this study, we investigated the genetic architecture of flavonoid-related metabolites using metabolomic data from 200 soybean accessions together with whole-genome sequence data available for 198 accessions. We applied principal component analysis (PCA)-based metabolome GWAS to identify key loci and subsequently quantified single-nucleotide polymorphism (SNP) effects in different genetic backgrounds by stratifying genotypes based on major loci. Through these analyses, we demonstrate that metabolic traits under strong genetic control are governed by a few loci exhibiting both additive effects and non-additive (epistatic) interactions, thereby clarifying a hierarchical and context-dependent genetic architecture of metabolic traits.

## Materials and Methods

### Plant materials and field trial

We used 200 soybean (*Glycine max*) accessions from the mini core collection registered in the NARO Genebank (National Agriculture and Food Research Organization, Tsukuba, Japan). Field trials were conducted over two years, 2017 and 2018, in sandy soil fields at the Arid Land Research Center, Tottori University, Japan (35°32′ N, 134°12′ E; 14 m above sea level). Two levels of irrigation treatment were established (irrigation and drought), with three individuals per plot. Across both treatment groups, 200 accessions were arranged in two rows, each serviced by a single irrigation line. Microplots were situated in parallel on both sides of these lines. Within each treatment, the accessions were randomly distributed, with three plants per accession grown in each microplot. Each row was covered with white mulching to prevent infiltration of rainwater. Plants in the irrigated treatment received irrigation for a total of 5 h per day, whereas no irrigation was applied in the drought treatment. Planting geometry followed a spacing of 50 cm between rows, 80 cm between microplots, and 30 cm between individual plants. The field was pre-fertilized before sowing with N, P, K, Mg, and Ca at rates of 1.3, 1.2, 4.0, 2.2, and 1.7 g m⁻², respectively. Seeds were sown in May of both 2017 and 2018. Initially, two to three seeds were sown at each site. At 2 weeks post-germination, seedlings were thinned so that only one seedling was retained.

### Whole-genome single-nucleotide polymorphism data

Whole-genome sequence data were available for 198 soybean accessions from a previous study (Kajiya-Kanegae et al., 2021). All accessions were genotyped using the Illumina HiSeq X Ten or HiSeq 4000 (Illumina). The variants for each accession were called using the GATK HaplotypeCaller (release 4.0.4.0) with the ‘.g.vcf’ extension. GATK GenomicsDBImport and GenotypeGVCFs were used for joint genotyping to produce a single VCF per sample of GVCF. Then, variants underwent quality assessment using the GATK best practices pipeline (https://software.broadinstitute.org/gatk/best-practices/; last accessed on 15 January 2021) to obtain a raw VCF that passed through the variant filtration step. The initial step for the variant dataset contained 10,116,707 SNPs and 2,835,680 indels. After removing the indels, heterozygous genotypes were replaced with missing data based on the assumption that all markers were fixed in all the accessions in the mini core collections. The minor allele frequency (MAF) ≥ 0.025 was set as an additional filter. The default parameters of Beagle 5.0 were used for the imputation of both datasets. Finally, 4,776,813 SNPs were identified. Genotypes at each SNP were coded as 0, 1, or 2, where homozygotes for the reference allele were coded as 0, heterozygotes as 1, and homozygotes for the alternative allele as 2.

### Metabolome analysis

At two months after sowing, three individuals were selected for each of the 200 accessions in each treatment. Samples were collected from the uppermost fully expanded leaves of each individual, immediately frozen, and stored at −80 °C until metabolome analysis. Targeted LC–MS/MS analysis was performed to quantify 188 metabolites (Supplementary Table 1).

Metabolome analysis was conducted using a previously published pipeline (Sawada et al., 2009; Uchida et al., 2020). Freeze-dried leaf samples (4 mg) were extracted with 1 mL of 80% (v/v) methanol containing 0.1% (v/v) formic acid, 8.4 nM lidocaine, and 210 nM 10-camphorsulfonic acid as internal standards. LC–MS/MS analysis was performed using a Nexera X2 UPLC system coupled to an LCMS-8050 triple quadrupole mass spectrometer (Shimadzu, Kyoto, Japan). The scheduled multiple reaction monitoring (MRM) transitions for each metabolite are listed in Supplementary Table 1. Raw data (.lcd files) were converted to Abf format using the Reifycs Abf Converter (https://www.reifycs.com/AbfConverter/), and peak areas were calculated using MRMPROBS (Tsugawa et al., 2013).

### Outlier removal

Outliers were removed from the metabolome data for each year. First, a regression analysis was performed for each metabolite, with the metabolite as the dependent variable and the genotype and treatment as explanatory variables. A linear mixed-effects model was used in which genotype was treated as a random effect and treatment as a fixed effect:

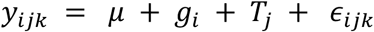

where:

*y_ijk_* is the observed metabolite value for the *i*-th genotype, under the *j*-th treatment, for the *k*-th replicate. *μ* is the overall mean (fixed intercept), *g_i_* is the random effect of the *i*-th genotype, *T_j_* is the fixed effect of the *j*-th treatment, and *ε_i_*_j*k*_ is the residual error. The random effects were assumed to be independent and identically normally distributed as follows: 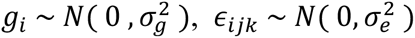

Based on this model, the best linear unbiased predictors (BLUPs) for each genotype were calculated for each metabolite, effectively removing the fixed treatment effects and residual error. Residual analyses were then performed, and observations with residuals outside the following range defined as outliers: mean residual ± 5 × standard deviation of the residuals. This procedure was applied to the combined population across both irrigation and drought treatments.

All analyses were conducted using the “lmer” function of the “lme4” package (Bates et al., 2015) in R (version 4.4.2; R Core Team 2024).

### Principal component analysis

PCA was performed for the metabolite data from the combined population of irrigation and drought treatments (200 accessions × 2 treatments × 3 individuals/accession/treatment = 1,200 individuals). Probabilistic PCA (PPCA) was used, which estimates the principal component (PC) scores while imputing missing values, including those from failed measurements or removed outliers. PCA was conducted separately for three metabolite datasets: all detected metabolites (188 metabolites), flavonoid-related metabolites (83 metabolites), and metabolites with high broad-sense heritability(ℎ^2^ > 0.9), which comprised 40 metabolites in 2017 and 43 metabolites in 2018. To focus on PCs that better capture data variance, we selected the top PCs until the cumulative contribution ratio reached approximately 0.5. For the 188 metabolites analyzed in 2017, this criterion resulted in the extraction of PC1 through PC6 (Supplementary Figure 1F). Accordingly, we consistently targeted PC1–PC6 for all other subsequent analyses. Factor loadings were calculated as the correlation coefficients between the original metabolite data and the PC scores, excluding sample pairs with missing values.

PCA was performed using the “pca” function from the “pcaMethods” package (Stacklies et al., 2007) in R. The arguments were set to method = “ppca” to specify PPCA and nPcs = 6 for the number of PCs. Factor loadings were calculated using the “cor” function with the argument use = “pairwise.complete.obs” to exclude pairs with missing values.

### Prediction of best linear unbiased predictors and estimation of broad-sense heritability

For each metabolite or PC, a linear mixed-effects model was fitted, with genotype as a random effect and treatment as a fixed effect using the “lmer” function in the R package “lme4”. BLUPs for each genotype were obtained from the fitted model after accounting for the fixed effect of treatment. Variance components for genotype and residual error were estimated from the fitted model.

Broad-sense heritability at the genotype level was estimated from the variance components obtained from the mixed-effects model as follows:

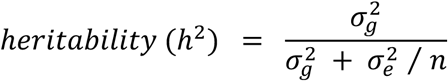

where:

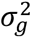 is the genetic variance, 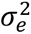 is the error variance, and *n* is the number of individual replicates per genotype.

### Genome-wide association study

For each metabolite or PC, GWAS was conducted using a linear mixed model. The BLUPs for each genotype were used as the dependent variable. The SNP marker being tested and population structure were included as fixed effects, and the polygenic background was modeled as a random effect. The model was specified as:

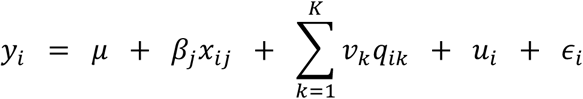

where:

*y_i_* is the BLUP for the *i*-th genotype, *μ* is the overall mean, *x_ij_* is the genotype value of the *j*-th SNP for the *i*-th genotype. *β_j_* is a fixed effect of the *j*-th SNP, *v_k_q_ik_* is a fixed effect for the *i*-th genotype of the *k*-th PC used to correct for population structure (in this case, the first two PCs served as covariates), *u_i_* is the random polygenic background effect, assumed to follow a multivariate normal distribution 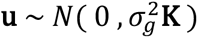, where K is the genomic relationship matrix (GRM), and *ε_i_* is the residual error, assumed to follow a multivariate normal distribution 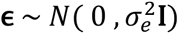.

BLUPs were obtained for each genotype based on individual metabolite values or PC scores. Population structure covariates were derived from the eigenvalue decomposition of GRM. A total of 405,409 SNP markers were used after filtering for minor allele frequency ≥ 0.05 and linkage disequilibrium ≤ 0.95. Statistical significance of SNP effects was evaluated using the Wald test. Multiple testing correction was performed using the Benjamini–Hochberg method (Benjamini and Hochberg, 1995), with a false discovery rate of 0.05. Manhattan plots were generated using −log_10_(P) values based on the adjusted *P* values.

GWAS analyses were conducted using 195 accessions for which both metabolomic and genomic data were available. For SNPs identified in GWAS of PCs, the population was stratified according to allelic combinations at these SNPs, and subsequent analyses were performed using the corresponding genotype subset. All GWAS analyses were performed using the “gaston” package (Perdry and Dandine-Roulland, 2023) in R. GRM was calculated from the SNP genotype data using the “GRM” function implemented in the “gaston” package. Model fitting and SNP testing were conducted using the “association.test” function, with the following arguments: method = “lmm”, response = “quantitative”, and test = “wald”. The number of PCs used to correct for population structure was set to p = 2.

### Estimation of single-nucleotide polymorphism regression coefficients in partitioned groups

Groups partitioned based on the allelic status of detected SNPs were used to estimate SNP regression coefficients for each metabolite via GWAS. Analyses were conducted using either the full set of 195 accessions or subgroups defined by single or combined SNP genotypes. For all detected SNPs, genotypes were homozygous (coded as 0 or 2). The regression model was identical to that used in the other GWAS analyses. To simplify the description of multilocus genotype groups throughout the manuscript, an asterisk (*) denotes an arbitrary genotype at the corresponding SNP, whereas specified positions indicate fixed genotypes.

### Construction of a graphical model using a Bayesian network

We constructed graphical models representing the network structure among SNPs and either metabolites or PCs using a Bayesian network framework. This framework represents relationships among nodes through conditional probabilities and estimates the graph structure using a score-based approach. We used the following as input data: (i) genotypes of four SNPs identified by GWAS of PCs and genotype-level BLUPs of 40 metabolites exhibiting high heritability; and (ii) genotypes of the same four SNPs and genotype-level BLUPs of the four PCs for which these SNPs were detected. Structure learning of the Bayesian network was performed using the “hc” function from the “bnlearn” package in R. The argument was set to score = “bic-cg”. No whitelist or blacklist constraints were specified, and neither the presence nor the direction of edges between variables was predefined. Thus, the network structure was inferred entirely from the data using a structure-learning algorithm.

### Partitioning of accessions by single-nucleotide polymorphism genotype and phylogenetic group

Based on whole-genome sequencing data, the soybean accessions used in this study were previously classified into three phylogenetic groups: Japan, World, and Primitive (Kajiya-Kanegae et al., 2021). Accessions were then partitioned according to the genotype at each SNP identified by GWAS of PCs or according to multilocus genotype combinations. Within each genotype-based subgroup, the number of accessions belonging to each phylogenetic group was calculated.

### Reference for metabolite and gene annotations and pathway information

Metabolic pathway information was obtained from the Kyoto Encyclopedia of Genes and Genomes (KEGG) database (https://www.genome.jp/kegg/). Gene information, including genomic locations and functional annotations, was retrieved from Phytozome (https://phytozome-next.jgi.doe.gov/) and KEGG. In this study, according to the KEGG PATHWAY database, metabolites involved in the shikimate pathway, phenylpropanoid pathway, flavonoid biosynthesis pathway, isoflavonoid biosynthesis pathway, flavone and flavonol biosynthesis pathway, anthocyanin biosynthesis pathway, and flavonoid metabolic pathway were defined as flavonoid-related metabolites.

## Results

### Broad-sense heritability

Field trials were conducted in 2017 and 2018, using 200 soybean accessions that were irrigated or exposed to drought conditions. Genotype-level broad-sense heritability was estimated for 188 metabolites. The proportion of metabolites with heritability values greater than 0.5 was 0.670 in 2017 and 0.793 in 2018 (Supplementary Figure 1A, E). In both years, heritability values exceeded 0.9 for many metabolites. These results indicate that many of these metabolites are under strong genetic control.

### GWAS of PCs for 188 metabolites and candidate gene identification

To summarize the 188 metabolite variables, PPCA was performed. The first six PCs (PC1–PC6) were retained based on the leveling-off of explained variance (Supplementary Figure 1F). The heritability of all PCs was moderate to high (>0.5; Supplementary Figure 1B). GWAS of PCs identified significant associations between chromosome 6 and PC1–PC3 (Supplementary Figure 2). Notably, PC2 showed a strong peak at Chr06_18760995 (−log_10_P > 50). Metabolites strongly associated with PC2 (|loading|≥0.6; Supplementary Table 2A, B) were predominantly involved in the flavonoid biosynthesis pathway (Supplementary Figure 3). GWAS of these metabolites identified peaks in genomic regions highly proximal to the peak for PC2, whereas similar peaks were not observed for metabolites associated with PC1 or PC3. Among the 14 candidate genes detected in the PC2-associated region (−log_10_P ≥20) (Supplementary Table 3), Glyma.06G202300 was annotated as flavonoid-3′-hydroxylase (F3′H), which is a key enzyme for flavonoid biosynthesis. Two metabolites (kaempferol and quercetin) corresponded to the substrate and product of the reaction catalyzed by this enzyme (Supplementary Figure 3).

Notably, the genotype at Chr06_18760995 showed the opposite effect on these metabolites. For kaempferol, genotype-level BLUPs increased and decreased in genotype 2 and 0, respectively, whereas the opposite trends were observed for quercetin (Supplementary Figure 4A, B). PC2 scores clearly separated genotypes at this SNP, whereas PC1 scores did not (Supplementary Figure 5A). Furthermore, the SNP was strongly associated with pubescence color (Toda et al., 2005). All accessions with tawny or intermediate pubescence belonged to genotype 0, while most accessions with gray pubescence belonged to genotype 2 (Supplementary Figure 6), consistent with the known function of F3′H. Hereafter, Chr06_18760995 is referred to as Chr06_18760995 (F3′H).

### GWAS of PCs for flavonoid-related metabolites and identification of additional significant loci

To further investigate genetic regions associated with flavonoid-related metabolites, we focused on 83 metabolites annotated to the flavonoid pathway. PCA followed by GWAS identified a strong peak on chromosome 6 for PC1 (−log_10_P > 50; Supplementary Figure 7), consistent with the results from the analysis of the full set of 188 metabolites (Supplementary Figures 2B, 7A). In addition, a distinct peak for PC4 was detected on chromosome 10, with the lead SNP at Chr10_42562665 (Supplementary Figure 7). All PCs showed relatively high heritability (*h*^2^ > 0.6; Supplementary Figure 1C).

Among the 188 metabolites, 40 exhibited exceptionally high heritability (ℎ^2^ ≥ 0.9), all of which were flavonoid-related. GWAS of PCs derived from these 40 metabolites identified strong peaks on chromosomes 6 and 10 (PC1 and PC3, Figure 1A, C), consistent with the results from the analysis of the other metabolite sets (Supplementary Figures 2B, 7A, 7D). Additional significant signals were detected on chromosome 6 (PC2) and on chromosome 17 (PC4), with the lead SNPs Chr06_47490224 and Chr17_16065902, respectively. These PCs also exhibited high heritability (ℎ^2^ > 0.7; Supplementary Figure 1D).

**Figure 1.**
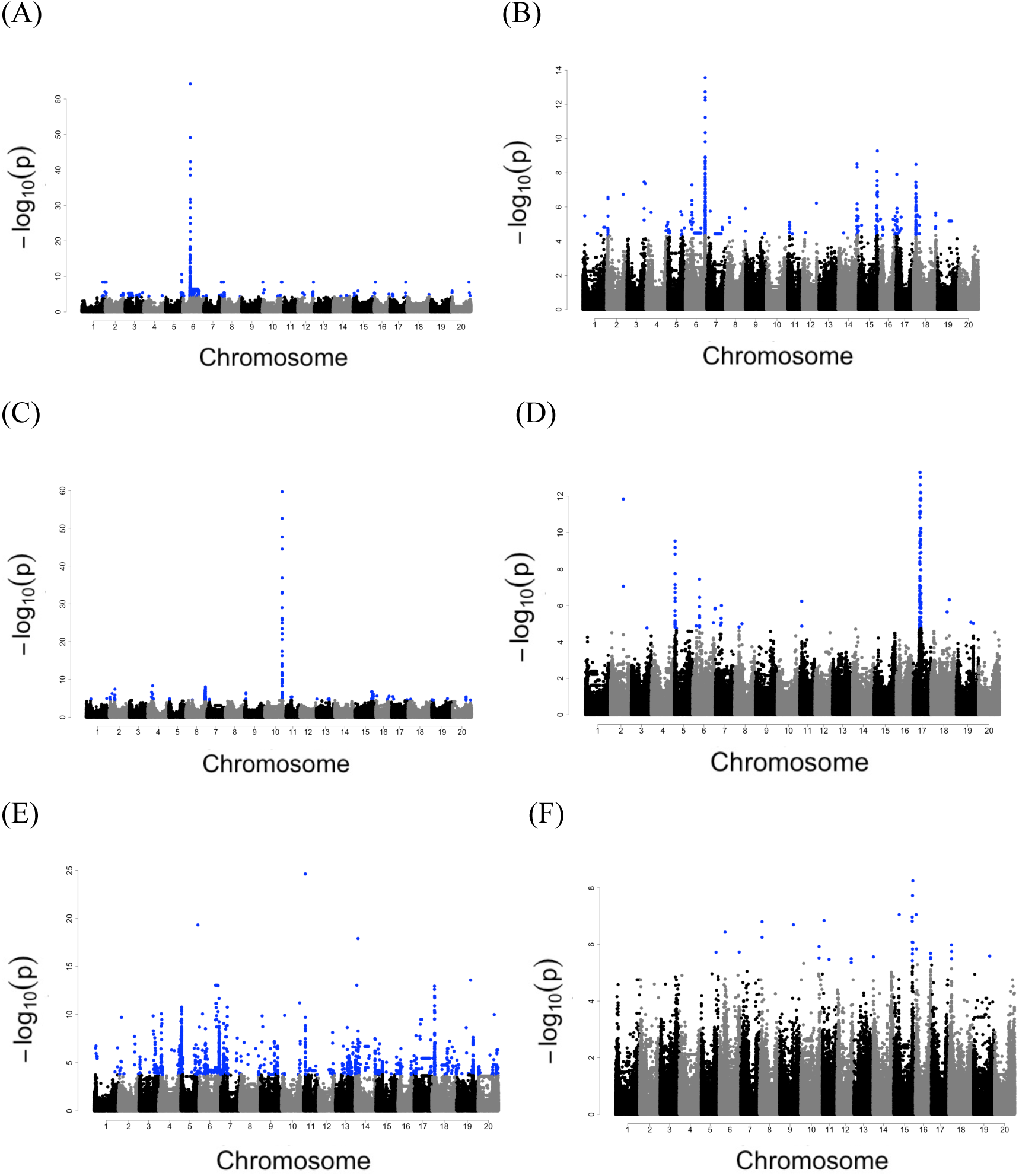
GWAS of PCs for the 40 metabolites with the highest heritability. Manhattan plots were produced based on GWAS results for (**A–F**) PC1–PC6. PCs were generated from 40 metabolites with an estimated broad-sense heritability of at least 0.9 in 2017. The horizontal axis presents the position on each chromosome, whereas the vertical axis presents the −log_10_(P) value. Blue dots denote SNPs that exceeded the significance threshold.

Analysis of 2018 data identified SNPs at the same or nearby genomic regions, confirming the robustness of these associations (Supplementary Figure 8).

### Effects of the detected SNPs on the flavonoid biosynthetic pathway

The four SNPs identified (Chr06_18760995 (F3′H), Chr06_47490224, Chr10_42562665, and Chr17_16065902) were associated with variation in the abundance of flavonoid-related metabolites. To evaluate their effects at the metabolite level, GWAS was performed for each of the 83 flavonoid-related metabolites. Among these metabolites, 28 showed significant associations near Chr06_18760995 (F3′H), while 7 and 5 were associated with Chr06_47490224 and Chr10_42562665, respectively. No significant associations were detected near Chr17_16065902. These results indicate that Chr06_18760995 (F3′H) has the strongest and most widespread effect on the flavonoid-related metabolites among the four SNPs.

### Estimation of other SNP effects in groups partitioned by the most effective SNP

To evaluate whether the effects of other SNPs depend on the genotype at Chr06_18760995 (F3′H), the 195 accessions were partitioned according to the genotype at this locus (0: n = 139; 2: n = 56), and SNP effects were estimated separately within each subgroup using the same regression model as that used for the GWAS.

For many flavonoid-related metabolites, the regression coefficients of the other three SNPs (Chr06_47490224, Chr10_42562665, and Chr17_16065902) were close to zero in the genotype 2 group, but substantially greater in the genotype 0 group (Figure 2A, E, J). This pattern was consistently observed for flavonoid-related metabolites, but not for non-flavonoid metabolites. For each SNP, the distribution of regression coefficients for non-flavonoid metabolites was used as an empirical null distribution, and the mean ± 3 SD of this distribution was used as the threshold for identifying metabolites with unusually large SNP effects. Using this criterion, 27, 28, and 7 metabolites were identified for Chr06_47490224, Chr10_42562665, and Chr17_16065902, respectively (Figure 2A, E, J). In most cases, SNP effects were detectable only in one of the Chr06_18760995 genotype groups. Analyses of genotype-level BLUPs across multilocus genotype combinations further showed that the effect size and direction of these SNPs depended on the Chr06_18760995 genotype (Figure 2B–D, F–I, K, L). These results indicate that the effects of Chr06_47490224, Chr10_42562665, and Chr17_16065902 on flavonoid-related metabolites are conditional on the genotype at Chr06_18760995 (F3′H).

**Figure 2.**
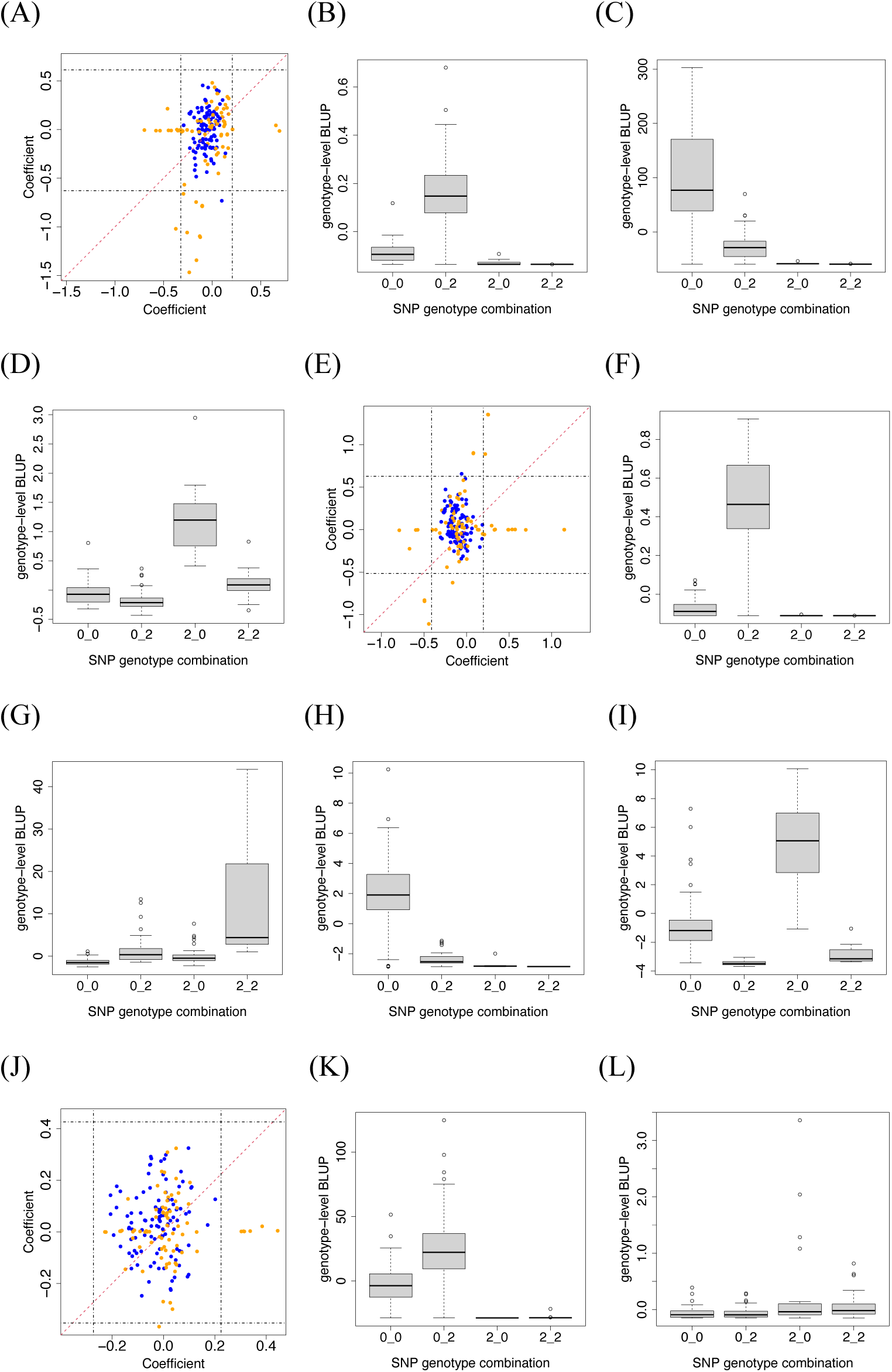
Estimated SNP regression coefficients and genotype-level BLUPs for metabolites. (**A, E, J**) Scatter plots of the regression coefficients for the SNPs (**A**) Chr06_47490224, (**E**) Chr10_42562665, and (**J**) Chr17_16065902 estimated for each metabolite. The horizontal axis presents coefficients from the population with genotype 0 of Chr06_18760995, whereas the vertical axis presents coefficients from the population with genotype 2 of Chr06_18760995. Each dot indicates a flavonoid-related (orange) or non-flavonoid (blue) metabolite. Vertical and horizontal dashed lines indicate the mean ± 3 SD calculated from the regression coefficients of non-flavonoid related metabolites. (**B–D, F–I, K, L**) Boxplots of genotype-level BLUPs in populations partitioned by two-locus SNP genotypes. The vertical axis presents BLUPs for each metabolite, predicted with a mean of 0. The horizontal axis presents the two-locus genotype combination, where the left number is the genotype of Chr06_18760995 and the right number is the genotype of (**B**–**D**) Chr06_47490224, (**F**–**I**) Chr10_42562665, or (**K**, **L**) Chr17_16065902. The following metabolites are those with the largest absolute regression coefficient within each of the four quadrants outside the mean ± 3 SD: (**B**) X00431, (**C**) X200087, (**D**) X00922, (**F**) X500081, (**G**) X500099, (**H**) X200086, (**I**) X01128, (**K**) X00419, and (**L**) X00414.

### Graphical models of SNP-metabolite relationships

To infer the relationships among the four SNPs and metabolites, graphical models were constructed using a Bayesian network framework. When PCs derived from the 40 metabolites were used, Chr06_18760995 (F3′H) was identified as the root node. Chr06_47490224 was positioned downstream of Chr06_18760995, while Chr10_42562665 and Chr17_16065902 were positioned further downstream of Chr06_47490224 (Figure 3A). A similar hierarchical structure was inferred when individual metabolites were used instead of PCs (Figure 3B). The numbers of metabolites assigned as child nodes were 27 for Chr06_18760995 (F3′H), 17 for Chr06_47490224, 21 for Chr10_42562665, and 6 for Chr17_16065902. Among the 27 metabolites assigned as child nodes of Chr06_18760995, 19 showed strong or moderate GWAS signals near this locus (Supplementary Table 4). Comparison with PCA loadings further supported the inferred SNP–metabolite relationships. For each SNP, the PC showing the strongest GWAS signal was identified, and the number of top-ranked metabolites based on the absolute values of their loadings was matched to the number of metabolites assigned as child nodes of that SNP in the Bayesian network. The overlap between the two metabolite sets was then evaluated. For example, 20 of the 27 metabolites (0.74) assigned as child nodes of Chr06_18760995 overlapped with the 27 top-ranked metabolites based on the absolute values of their loadings for PC1. The corresponding overlap proportions were 0.47, 0.71, and 0.33 for Chr06_47490224, Chr10_42562665, and Chr17_16065902, respectively (Supplementary Table 4).

**Figure 3.**
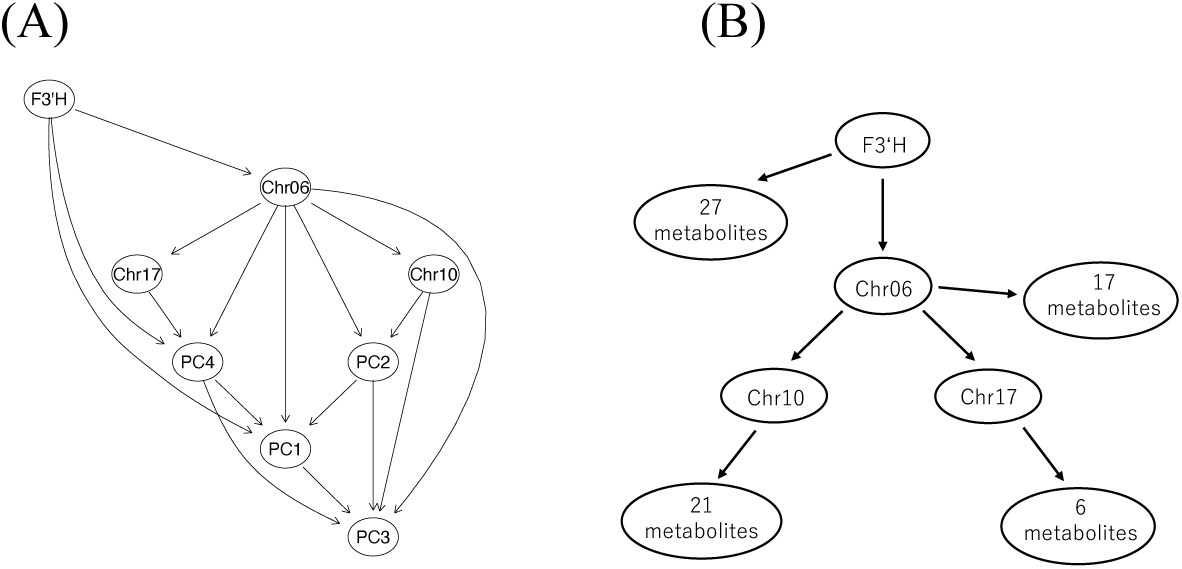
Bayesian network structures inferred from SNP genotypes and either PCs or highly heritable metabolites. Bayesian network structures inferred from (**A**) the PCs for 40 highly heritable metabolites and (**B**) the 40 highly heritable metabolites together with the genotypes of the four SNPs identified by GWAS of PCs. Arrows indicate directed edges inferred between variables to maximize the overall network score. In graphs, F3′H, Chr06, Chr10, and Chr17 represent the SNPs Chr06_18760995, Chr06_47490224, Chr10_42562665, and Chr17_16065902, respectively. In (**B**), for clarity, the child metabolites of each SNP are represented as a single node labeled with the number of metabolites assigned to that SNP. Considering that a metabolite can be assigned as a child node of more than one SNP, these groups are not mutually exclusive.

Together, these results support a hierarchical SNP-metabolite dependency structure in which Chr06_18760995 (F3′H) and Chr06_47490224 are positioned upstream of Chr10_42562665 and Chr17_16065902.

### Estimation of downstream SNP effects under multilocus genetic backgrounds

On the basis of the hierarchical relationships identified in the Bayesian network, we evaluated whether the effects of downstream SNPs depend on multilocus genetic backgrounds defined by upstream SNPs. The 195 accessions were partitioned into four groups according to genotypes at Chr06_18760995 (F3′H) and Chr06_47490224: (0_0), (0_2), (2_0), and (2_2). Within each subgroup, GWAS was performed and regression coefficients were estimated for Chr10_42562665 and Chr17_16065902. The number of metabolites with significant effects (based on the 3 SD threshold) differed across subgroups. For Chr10_42562665, 22 and 7 metabolites were detected in the (0_*) and (2_*) genetic backgrounds, respectively. For Chr17_16065902, 4 and 3 metabolites were detected in the (0_*) and (2_*) genetic backgrounds, respectively (Figure 4A, E, G, J). Further analyses using three-locus genotype combinations confirmed that the magnitude and direction of SNP effects varied depending on the upstream genotype combinations (Figure 4B–D, F, H, I, K). These results indicate that the effects of Chr10_42562665 and Chr17_16065902 on flavonoid-related metabolites depend on the multilocus genetic background defined by Chr06_18760995 (F3′H) and Chr06_47490224.

**Figure 4.**
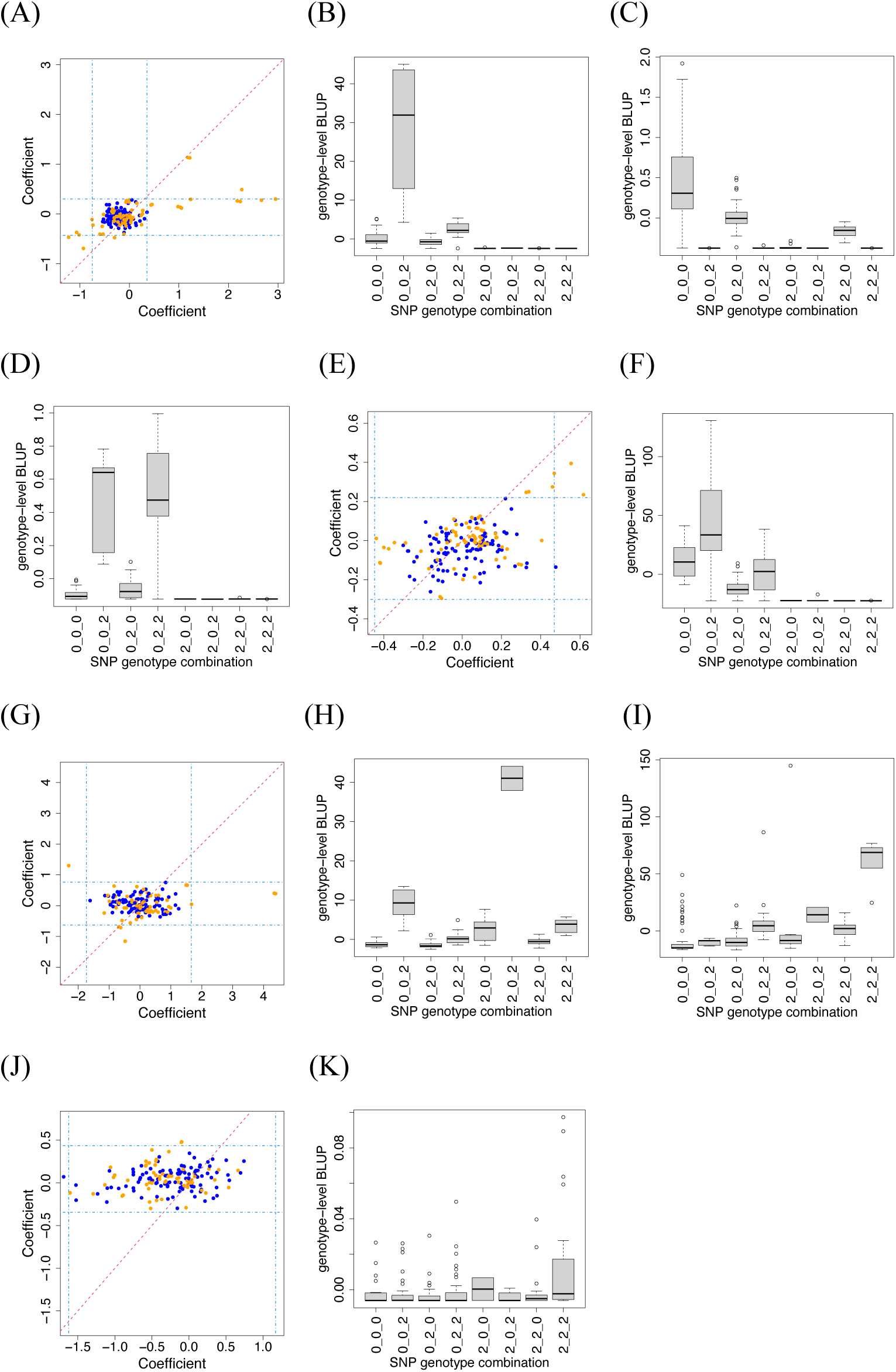
Estimated SNP regression coefficients and genotype-level BLUPs for each metabolite. (**A, E, G, J**) Scatter plots of SNP regression coefficients for each metabolite. For (**A**) Chr10_42562665 and (**E**) Chr17_16065902, the horizontal axis presents coefficients estimated in the population with the (Chr06_18760995_Chr06_47490224) = (0_0) genotype combination, whereas the vertical axis presents those from the (0_2) population. For (**G**) Chr10_42562665 and (**J**) Chr17_16065902, the axes present coefficients from the (2_0) and (2_2) populations, respectively. Each dot indicates a flavonoid-related (orange) or non-flavonoid (blue) metabolite. Vertical and horizontal dashed lines indicate the mean ± 3 SD calculated from the regression coefficients of the non-flavonoid metabolites. (**B–D, F, H, I, K**) Boxplots of genotype-level BLUPs in populations partitioned by three-locus SNP genotype combinations. The vertical axis presents BLUPs for each metabolite. The horizontal axis presents the three-locus genotype combination: (**B–D**) (Chr06_18760995_Chr06_47490224_Chr10_42562665), and (**F, H, I, K**) (Chr06_18760995_Chr06_47490224_Chr17_16065902). The following metabolites are the flavonoid-related metabolites with the largest absolute regression coefficient within each of the four quadrants outside the mean ± 3 SD: (**B**) X500098, (**C**) X01174, (**D**) X01219, (**F**) X200004, (**H**) X500099, **(I)** X00853, and (**K**) X400001.

### Association between phylogenetic groups and SNP genotypes

A previous study classified accessions into three phylogenetic groups (Japan, World, and Primitive) on the basis of whole-genome sequencing data (Kajiya-Kanegae et al., 2021). Using this classification, we examined the distribution of these three groups across SNP genotypes and their combinations. Hereafter, multilocus genotypes are represented as four-digit genotype combinations with the order of SNPs defined in the legend of Figure 5.

**Figure 5.**
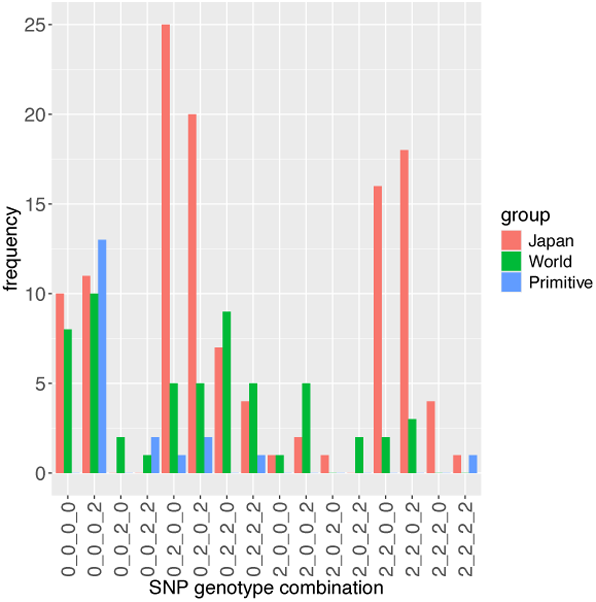
Number of genotypes in three phylogenetic groups according to four-locus genotype combinations. For 195 genotypes, the population was partitioned by the genotype combinations of four SNPs detected by GWAS of PCs. The number of genotypes belonging to each of three phylogenetic groups (Japan, World, and Primitive) within each partitioned group is presented. The following SNPs were used: Chr06_18760995, Chr06_47490224, Chr10_42562665, and Chr17_16065902. The horizontal axis presents the four-locus genotype combinations, with genotypes arranged in the order of the SNPs listed above. The vertical axis presents the number of genotypes.

Partitioning by individual SNPs revealed imbalanced genotype frequencies for Chr06_18760995 (F3′H), Chr06_47490224, and Chr10_42562665 (Figure 5). Notably, genotype 2 of Chr06_47490224 and genotype 0 of Chr10_42562665 were enriched in the Japan group, genotype 0 of Chr06_18760995 (F3′H), genotype 0 of Chr06_47490224, and genotype 2 of Chr17_16065902 were enriched in the Primitive group. When considering multilocus genotypes, clear patterns were detected for specific combinations. In particular, the (0_0_0_2) genotype was strongly enriched in Primitive accessions, whereas other combinations, such as (*_2_0_*), were enriched in Japan accessions (Figure 5). These results indicate that the genotypes of the four SNPs are associated with phylogenetic structure; they may be particularly useful for distinguishing between Japan and Primitive groups.

## Discussion

### Identification of genomic regions associated with metabolites and metabolic pathways

A key objective of this study was to identify genomic regions underlying variation in metabolite abundance. By completing PCA of high dimensional metabolomic data, we detected four major loci associated with flavonoid-related variation: Chr06_18760995 (F3′H), Chr06_47490224, Chr10_42562665, and Chr17_16065902. These loci were consistently detected across analyses involving 188 metabolites, 83 flavonoid-related metabolites, and a subset of 40 highly heritable metabolites. This consistency suggests that PCA effectively captured pathway-level metabolic variation, while reducing noise from metabolites under relatively weak genetic control. Notably, all loci detected in the large metabolite sets were also identified in the analysis of 40 highly heritable metabolites, all of which were related to flavonoids. These results suggest that the detected loci represent major genetic determinants of flavonoid-related metabolic variation. The consistent detection of these loci in both 2017 and 2018 further supports the stability of their effects.

The magnitude of SNP effects differed among loci. Chr06_18760995 (F3′H), Chr06_47490224, and Chr10_42562665 were clearly associated with individual metabolites, whereas Chr17_16065902 was not detected by a single-metabolite GWAS. The observed associations between the former loci and multiple metabolites likely reflect pleiotropy (Chen et al., 2014; Luo, 2015; Soltis and Kliebenstein, 2015). This suggests that the former loci directly and strongly affect metabolite variation, whereas Chr17_16065902 may have more subtle or coordinated effects detectable only through multivariate approaches (e.g., PCA).

### Hierarchical genetic control of context-dependent SNP effects

Among the four loci identified by PC-based GWAS, Chr06_18760995 had the strongest and most widespread effects on flavonoid-related metabolites. This locus was also associated with pubescence color, whose causal gene, *F3′ H*, encodes a key enzyme in the flavonoid pathway (Toda et al., 2005). Together with the strong GWAS signal, this suggests that the detected association reflects the effect of *F3′ H* or a variant in strong linkage disequilibrium with it.

Conditional analyses revealed that the effects of the other SNPs (Chr06_47490224, Chr10_42562665, and Chr17_16065902) depended on the genotype at Chr06_18760995. In particular, SNP effects were often attenuated or absent in one genotype background but detectable in another, reflecting strong context-dependent (epistatic) effects. Similar patterns were observed when considering multilocus combinations of Chr06_18760995 and Chr06_47490224, further supporting higher-order interactions among these loci. Although a few non-flavonoid metabolites were detected as significant, this was likely due to background variation or noise. In some cases, however, such metabolites may reflect unannotated or soybean-specific flavonoid-related processes.

The dependency of SNP effects is consistent with the hierarchical organization of metabolic pathways, where upstream genetic factors modulate downstream effects (Rowe et al., 2008). Bayesian network analyses further supported this interpretation, with Chr06_18760995 (F3′H) and Chr06_47490224 positioned upstream of Chr10_42562665 and Chr17_16065902 in the inferred network (Figure 3A, B). Notably, when variation at an upstream locus was associated with reduced pathway-level variation, the effects of downstream loci became largely undetectable regardless of their contribution to metabolite variation. By stratifying the population according to major upstream loci, we revealed such conditional effects and inferred hierarchical genetic control within the flavonoid biosynthetic pathway.

### Complex genetic control of highly-heritable metabolic networks

In multiple plant species, metabolites, particularly secondary metabolites, tend to exhibit relatively high heritability (Chen et al., 2014; Alseekh et al., 2015; Soltis and Kliebenstein, 2015). Moderate-to-high heritability (ℎ^2^ ≥ 0.5) was detected for a large proportion of the 188 metabolites analyzed in the current study, indicating strong genetic control. Notably, all 40 metabolites with exceptionally high heritability (ℎ^2^ ≥ 0.9) were related to flavonoids, suggesting that tightly regulated metabolites in this population are predominantly involved in flavonoid biosynthesis.

GWAS of PCs for these metabolites consistently identified four loci (Figure 1), whose effects were detectable across multiple scales, including PCs, individual metabolites, and subgroups of SNP genotypes. This indicates that these loci contribute to flavonoid variation at least partly through additive genetic effects. In addition, conditional analyses revealed strong context-dependent effects. The estimated effects of downstream SNPs varied markedly depending on the genotype at upstream loci, particularly Chr06_18760995 (F3′H) (Figures 2, 4). These patterns were consistently observed across metabolites and genetic backgrounds, providing strong evidence consistent with epistatic interactions among the four loci. Considered together, these results indicate that even highly heritable metabolites are influenced by a combination of additive and non-additive genetic effects, reflecting the complex genetic architecture underlying flavonoid metabolism.

### Diversity in genetic control patterns in a pathway

Previous metabolome GWAS research largely focused on pathway-level metabolite variations derived from additive genetic effects; however, similar investigations accounting for epistatic interactions remain unexplored (Chen et al., 2014; Wen et al., 2014; Luo, 2015). Our results address this gap by revealing that flavonoid-related metabolites under strong genetic control exhibited diverse patterns of genetic effects across the four loci. SNP effects varied considerably in both magnitude and direction depending on the genetic background, indicating that a given SNP can increase or decrease metabolite levels depending on the allelic context (Figures 2, 4). Notably, Chr06_18760995 (F3′H) had a strong negative effect in one genotype background, with similarly low metabolite levels, whereas substantial variations were retained in the alternative genetic background. By contrast, Chr06_47490224 had more moderate effects, indicating differences in allelic strength among loci. These observations indicate that the four loci play distinct genetic roles within the flavonoid pathway, differing in both the magnitude and context dependency of their effects. Despite this complexity, most of the observed variation could be explained by only four loci, suggesting that a few interacting genetic factors can generate substantial metabolic diversity.

### Association between SNP genotypes and phylogenetic structure

Considering that the four loci identified in this study clearly differed in terms of allele frequencies among phylogenetic groups, we examined whether the inferred pathway architecture was associated with the soybean population structure. The multilocus genotype (0_0_0_2) was enriched in Primitive accessions, whereas genotype 2 at Chr06_47490224 was enriched in Japan accessions, especially (*_2_0_*) (Figure 5). These patterns are consistent with the possibility that the genotype combination (0_0_0_2) represents an ancestral state, while genotype 2 at Chr06_47490224 is characteristic of the Japan group. Given that Chr06_47490224 significantly affected multiple flavonoid-related metabolites in the conditional GWAS (Figures 2E, 4A, 4G), this locus may contribute to lineage-associated variations in flavonoid metabolism in soybean. Flavonoid-related metabolites reportedly influence disease resistance, UV tolerance, and antioxidant activity (Agati et al., 2012; Falcone Ferreyra et al., 2012; Shen et al., 2022). The observed differences in allele frequencies of SNPs among phylogenetic groups may contribute to the variation in these physiological traits across different accessions or groups.

## Conclusion

In this study, we conducted a metabolome GWAS focusing on highly heritable flavonoid-related metabolites in soybean to elucidate their genetic control. Our results indicate that variation in these metabolites can largely be explained by only a few key loci acting through both additive and non-additive (epistatic) effects. Together, these findings reveal a hierarchical and context-dependent genetic architecture underlying flavonoid metabolic diversity in soybean.

## Author Contributions

TH: Data curation; Formal analysis; Investigation; Methodology; Software; Visualization; Writing – original draft.

KH: Software; Formal analysis; Methodology; Writing – review & editing.

YF: Data curation; Funding acquisition; Investigation; Writing – review & editing.

YT: Data curation; Funding acquisition; Investigation; Writing – review & editing.

YI: Funding acquisition; Investigation; Writing – review & editing.

YO: Funding acquisition; Investigation; Writing – review & editing.

YY: Funding acquisition; Investigation; Writing – review & editing.

HirT: Funding acquisition; Investigation; Writing – review & editing.

HidT: Funding acquisition; Investigation; Writing – review & editing.

MT: Funding acquisition; Investigation; Writing – review & editing.

HT: Funding acquisition; Investigation; Writing – review & editing.

AK: Funding acquisition; Investigation; Resources; Writing – review & editing.

MN: Funding acquisition; Investigation; Writing – review & editing.

TF: Funding acquisition; Investigation; Writing – review & editing.

MYH: Funding acquisition; Investigation; Writing – review & editing.

HI: Conceptualization; Funding acquisition; Investigation; Project administration; Supervision; Writing – review & editing.

## Acknowledgments

We are grateful to the technical staff at the Arid Land Research Center, Tottori University, and Ms. Izumi Higashida. We thank Dr. Ryota Yuasa (Chiba University, Japan) for valuable discussions and advice on the statistical and mathematical foundations of PCA, including matrix decomposition, decorrelation, orthogonal basis construction, and subspace representation, and Dr. Hideto Mochizuki for showing how to use the “bnlearn” package in R.

## Generative AI statement

The authors used generative AI to assist with refining the English language and improving the clarity of the manuscript. All scientific content, interpretations, and conclusions were developed and verified by the authors, who take full responsibility for the final version of the manuscript.

