## Supplementary file for "Metabolome genome-wide association study reveals hierarchical and epistatic genetic control of flavonoid metabolism in soybean"

**Supplementary Table 2. Metabolites extracted based on loadings for PC2**

IDs and annotations are presented for metabolites with an absolute loading of 0.6 or higher for PC2 generated using 2017 metabolite data. The listed metabolites had loadings of (**A**) $\geq 0.6$ and (**B**) $\leq -0.6$.

(A)

| **Sample ID** | **Name** |
|---|---|
| X00432 | Hyperoside |
| X200014 | 2i_Quercetin-3-arabinoside_quercetin-3-D-xyloside |
| X250001 | Quercetin 3-glucoside <Isoquercitrin> |
| X500080 | Quercetin 4′-glucoside |

(B)

| **Sample ID** | **Name** |
|---|---|
| X00422 | Kaempferol-3-O-glucoside |
| X00425 | Homoorietin |
| X00854 | Luteolin-4′-O-glucoside |
| X01128 | Cyanidin-3-O-rhamnoside chloride |
| X200012 | 2i_Maritimein_luteolin-7-O-glucoside |
| X210010 | Cyanidin |
| X500134 | Orobol |
| X210004 | Luteolin |

**Supplementary Table 3. Genes in the vicinity of Chr06_18760995**

The listed genes were located near the prominent peak detected by GWAS for PC2, generated from 188 metabolites using 2017 data. Genes located in the region with a −log10(P) value of 20 or higher were extracted and their functionally annotated.

| **Gene ID** | **Annotation** |
|---|---|
| 06G202300 | Flavonoid 3′-monooxygenase(hydroxylase) |
| 06G202400 | |
| 06G202500 | Activating signal cointegrator complex subunit 3 |
| 06G202600 | |
| 06G202700 | |
| 06G202800 | |
| 06G202900 | |
| 06G203000 | Glycosyl hydrolases family 28 |
| 06G203100 | |
| 06G203200 | F-type $H^+$-transporting ATPase subunit gamma |
| 06G203300 | Phosphoserine phosphatase |
| 06G203400 | Glutaminyl-tRNA synthetase |
| 06G203500 | Endonuclease/exonuclease/phosphatase family |
| 06G203600 | |

**Supplementary Table 4. Number of metabolites identified as child nodes or with high absolute loadings.**

The number of the metabolites inferred as child nodes for each of the four SNPs in the Bayesian network is presented. The same number of metabolites with the highest loadings on PCs corresponding to each SNP was extracted for comparisons. The counts for each set and the proportion of overlapping metabolites between the two sets for each SNP are presented.

| | **Child node** | **Highest absolute loadings** | **Overlap** | **Proportion** |
|---|---|---|---|---|
| Chr06_18760995 (PC1) | 27 | 27 | 20 | 0.74 (20/27) |
| Chr06_47490224 (PC2) | 17 | 17 | 8 | 0.47 (8/17) |
| Chr10_42562665 (PC3) | 21 | 21 | 15 | 0.71 (15/21) |
| Chr17_16065902 (PC4) | 6 | 6 | 2 | 0.33 (2/6) |

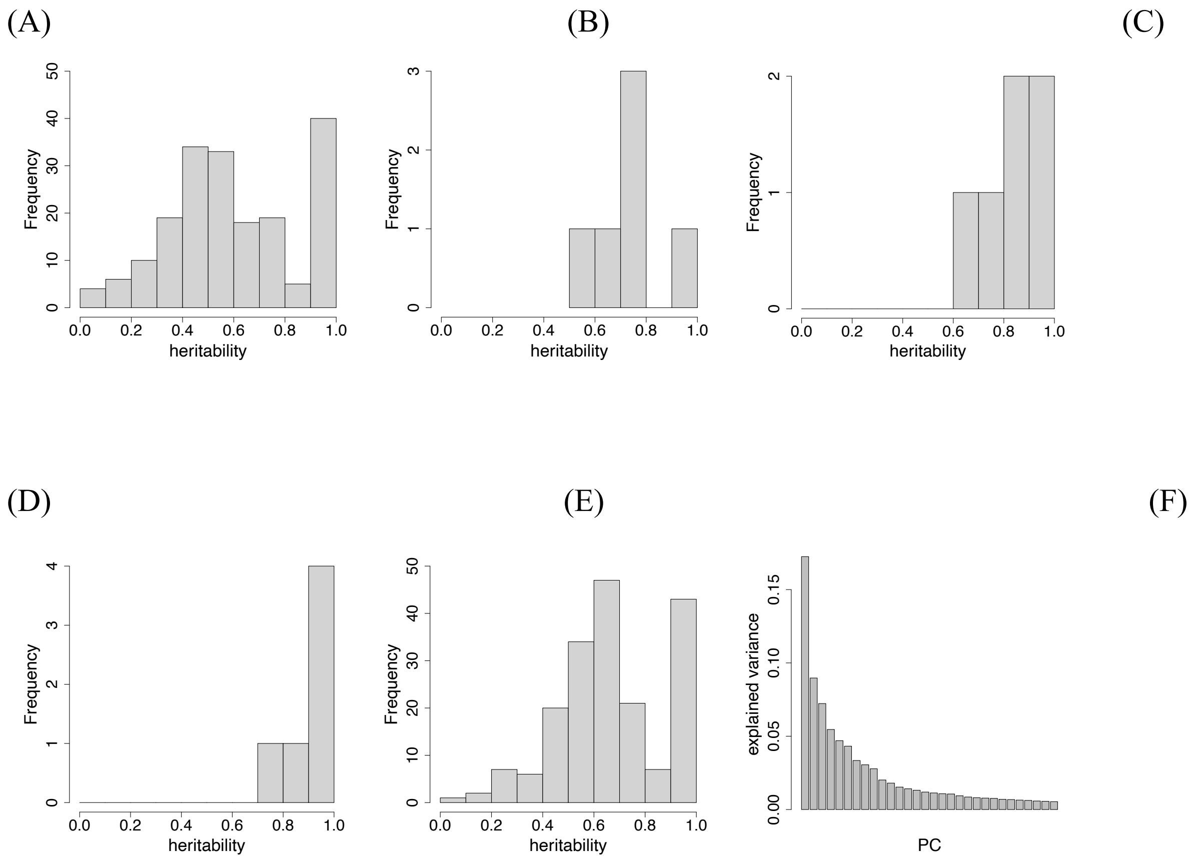


**Supplementary Figure 1. Estimated broad-sense heritability and explained variances of PCs.**

Histograms are presented for the estimated broad-sense heritability at the genotype level in 2017 for (**A**) all 188 metabolites, (**B**) PCs for the 188 metabolites, (**C**) PCs for 83 flavonoid-related metabolites, and (**D**) PCs for 40 metabolites with a heritability of 0.9 or higher. (**E**) Histogram of the estimated broad-sense heritability at the genotype level for 188 metabolites in 2018. The horizontal axis presents broad-sense heritability, whereas the vertical axis presents frequency. (**F**) Contribution ratio of each PC for the 188 metabolites in 2017.

(A)

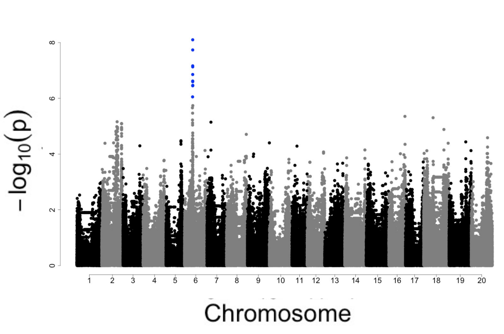


(B)

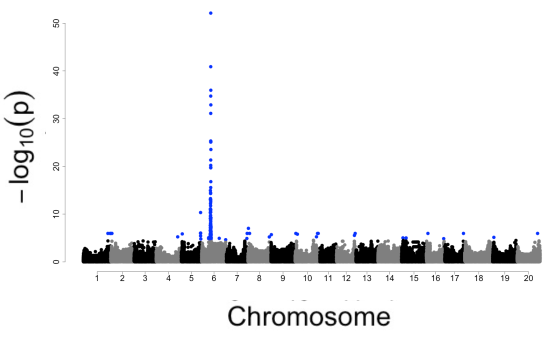


(C)

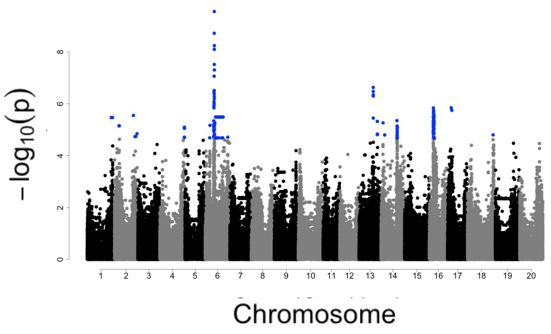


(D)

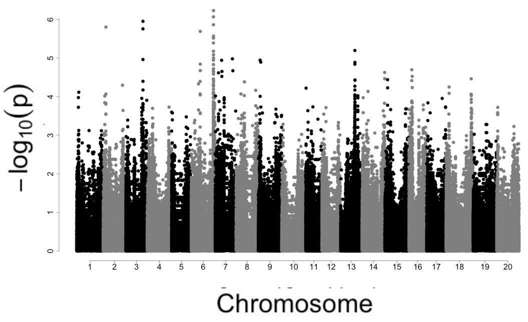


(E)

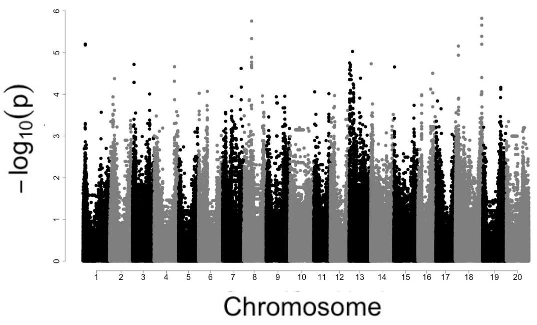


(F)

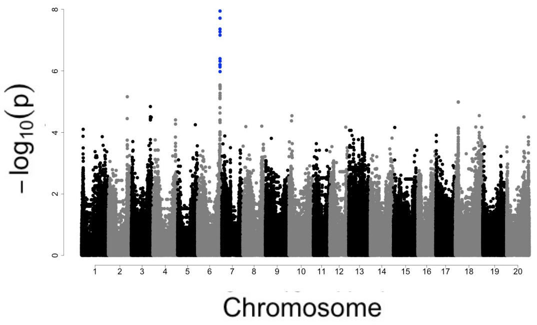


**Supplementary Figure 2. GWAS of PCs for 188 metabolites.**

Manhattan plots were produced based on GWAS results for (**A**–**F**) PC1–PC6, for 188 metabolites in 2017. The horizontal axis presents the position on each chromosome, whereas the vertical axis presents $-\log_{10}(P)$ values. Blue dots indicate SNPs that exceeded the significance threshold.

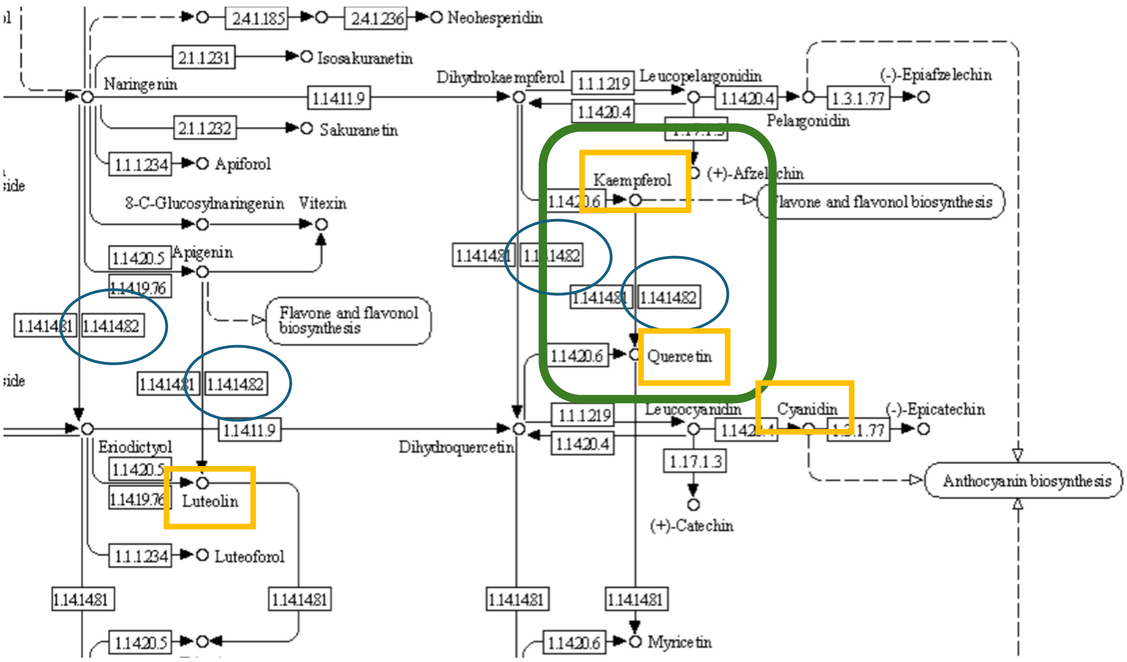


**Supplementary Figure 3. Mapping of metabolites with high absolute loadings on a metabolic pathway.**

The KEGG PATHWAY-derived figure presents the positions of metabolites in part of the flavonoid metabolic pathway. These metabolites were extracted according to their loadings for PC2, which was generated from the 188 metabolites using 2017 data. Yellow squares indicate mapped metabolites, while blue circles indicate *flavonoid 3'-monooxygenase* (*hydroxylase*). The metabolic reaction of interest (see main text) is outlined in green.

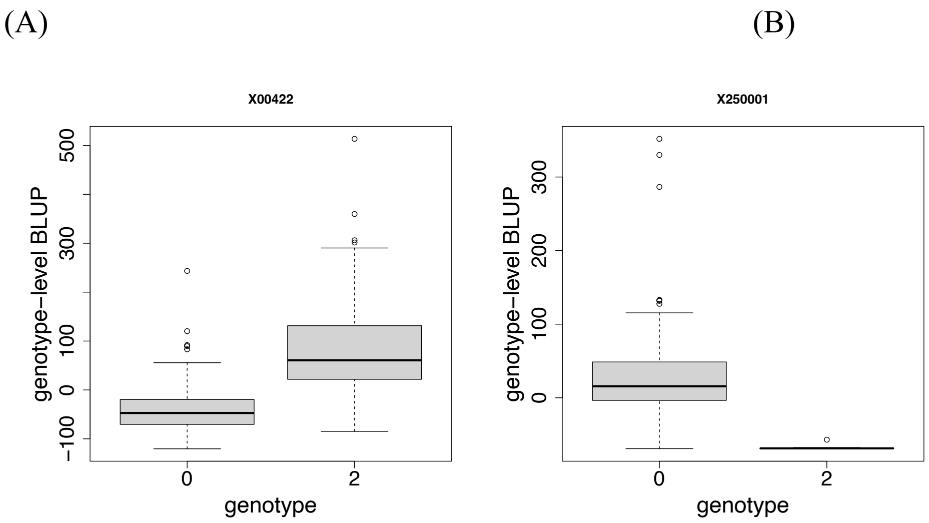


**Supplementary Figure 4. Genotype-level BLUPs for metabolites based on the genotype of Chr06_18760995.**

Boxplots present BLUPs for the derivative of (**A**) kaempferol (X00422) and (**B**) quercetin (X250001) in populations partitioned according to the genotype of Chr06_18760995.

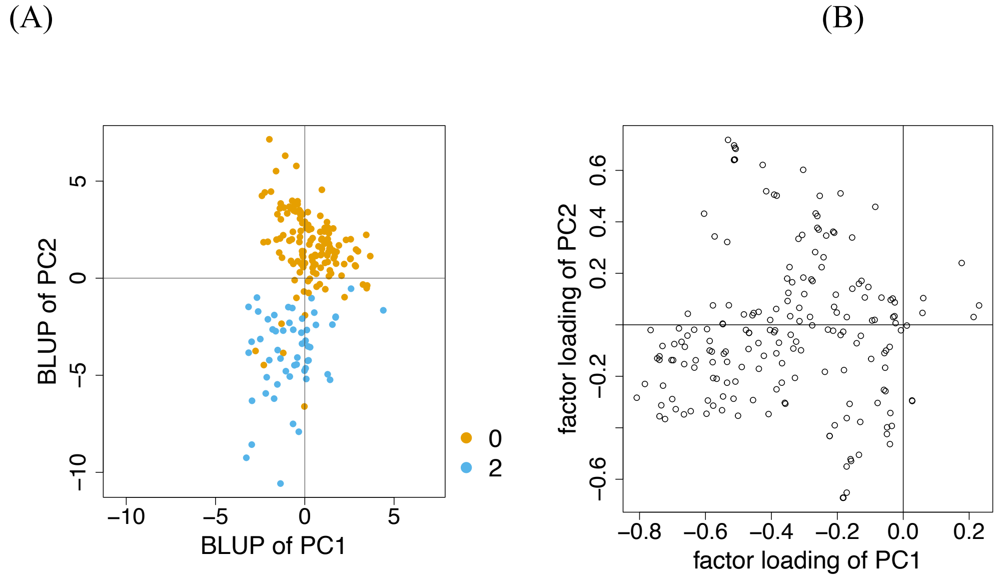


**Supplementary Figure 5. BLUPs of PCs for 188 metabolites and loadings.**

(**A**) Genotype-level BLUPs of PC1 and PC2 for 195 genotypes. Each dot represents a genotype. Genotypes are color-coded based on the allelic status of Chr06_18760995: orange for genotype 0 and blue for genotype 2. (**B**) Loadings on PC1 and PC2 for 188 metabolites. Each dot represents a metabolite. Both plots were generated using 2017 data.

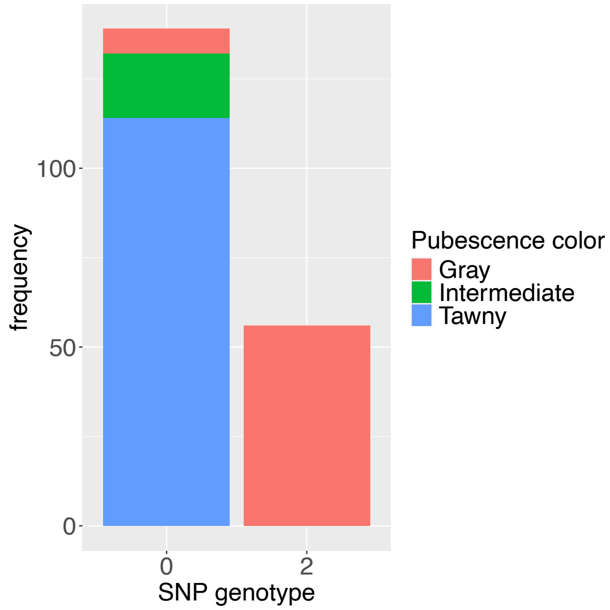


**Supplementary Figure 6. Pubescence color phenotypes and Chr06_18760995 genotypes.**

The number of genotypes for each trichome color phenotype is indicated for populations partitioned according to the genotype of Chr06_18760995.

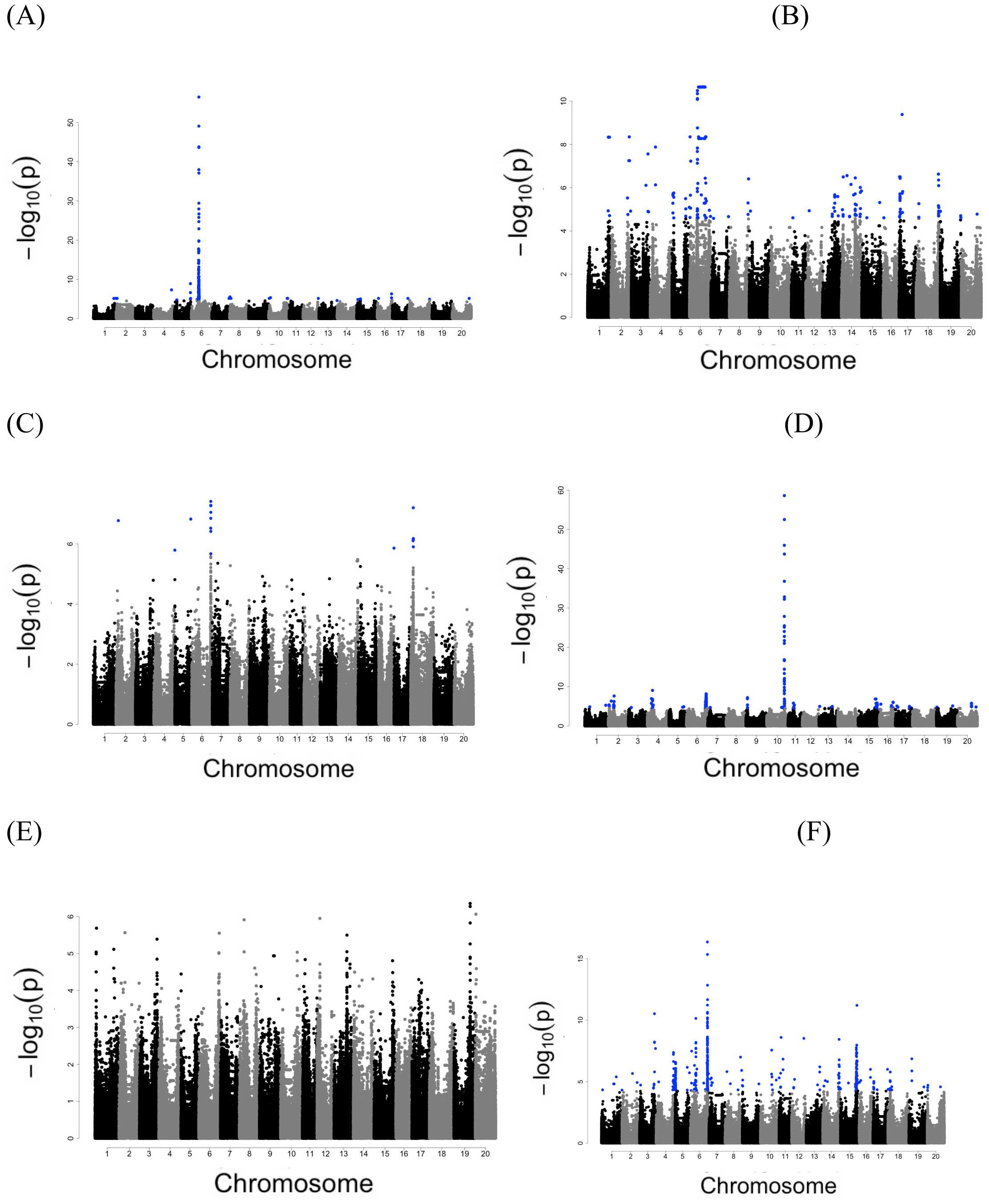


**Supplementary Figure 7. GWAS of PCs for metabolites of a specific metabolic pathway.**

Manhattan plots were produced based on GWAS results for (**A**–**F**) PC1–PC6, for 83 flavonoid-related metabolites in 2017. The horizontal axis presents the position on each chromosome, whereas the vertical axis presents $-\log_{10}(P)$ values. Blue dots indicate SNPs that exceeded the significance threshold.

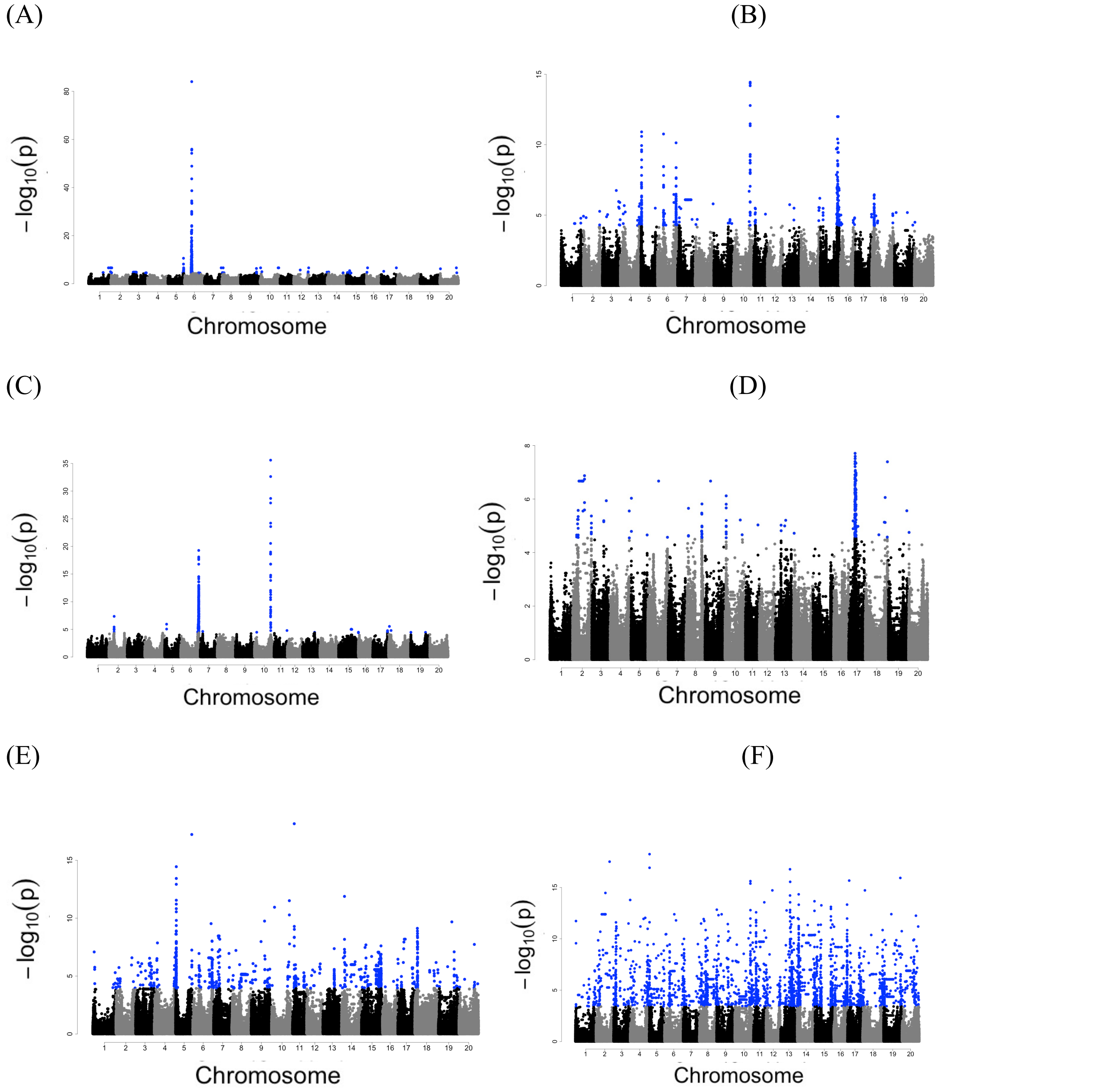


**Supplementary Figure 8. GWAS of PCs for highly heritable metabolites.**

Manhattan plots were produced based on GWAS results for (**A**–**F**) PC1–PC6, for metabolites with an estimated heritability of 0.9 or higher in 2018. The horizontal axis presents the position on each chromosome, whereas the vertical axis presents $-\log_{10}(P)$ values. Blue dots indicate SNPs that exceeded the significance threshold.
